# CerebrA: Accurate registration and manual label correction of Mindboggle-101 atlas for MNI-ICBM152 template

**DOI:** 10.1101/2019.12.19.883330

**Authors:** Ana L. Manera, Mahsa Dadar, Vladimir Fonov, D. Louis Collins

**Author notes:** Corresponding author(s): Louis Collins.

## Abstract

Accurate anatomical atlases are recognized as important tools in brain-imaging research. They are widely used to estimate disease-specific changes and therefore, are of great relevance in extracting regional information on volumetric variations in clinical cohorts in comparison to healthy populations. The use of high spatial resolution magnetic resonance imaging and the improvement in data preprocessing methods have enabled the study of structural volume changes on a wide range of disorders, particularly in neurodegenerative diseases where different brain morphometry analyses are being broadly used in an effort to improve diagnostic biomarkers.

In the present dataset, we introduce the Cerebrum Atlas (CerebrA) along with the MNI-ICBM2009c average template. MNI-ICBM2009c is the most recent version of the MNI-ICBM152 brain average, providing a higher level of anatomical details. Cerebra is based on an accurate non-linear registration of cortical and subcortical labelling from Mindboggle 101 to the symmetric MNI-ICBM2009c atlas, followed by manual editing.

## Background and Summary

Brain atlases are widely recognized as important tools in research for the analysis of neuroimages. High spatial resolution magnetic resonance imaging (MRI) and improved data preprocessing have enabled the study of structural volume changes on a wide range of disorders. Anatomical atlases, often averaged from multiple subjects, used to address disease-specific changes are crucial in order to provide regional information on volumetric variations in clinical cohorts in comparison to healthy populations.

The MNI-ICBM152 brain template2, from the Montreal Neurological Institute (MNI) is a crucial tool in neuroimage analysis. This multi-contrast atlas including T1w, T2w and PDw contrasts, was built recruiting brain scans from 152 young adults at 1.5 T. The 2009 edition uses group-wise non-linear registration for better alignment of cortical structures between subjects. The MNI-ICBM152 non-linear model has many advantages. It was created from a large number of subjects; hence it represents the average anatomy of the population and is not biased unlike single-subject models. In addition, the left-right symmetric version enables interpretation of asymmetries that might be found in an analysis.

Mindboggle-101 is the largest, publicly available set of manually labelled human brain images created from 101 human scans, labelled according to a surface-based cortical labelling protocol (DKT-Desikan-Killiany-Tourville labelling protocol)^1,3^. For the creation of the Mindboggle-101 dataset, developed to serve as brain atlas for use in labelling other brains, 101 T1-weighted (T1w) brain MRI images were selected and segmented based on a modification of the DKT cortical parcellation atlas^3^. These labels were then manually edited in agreement with the DKT protocol. Labelling was performed on the surface, yet, topographical landmarks visible in the folded surface were used to infer label boundaries. In addition, Mindboggle used non-cortical labels that were converted from Neuromorphometrics BrainCOLOR subcortex labels^4^.

The Cerebrum Atlas (CerebrA) includes co-registration of the Mindboggle atlas^1^ to the symmetric version of MNI-ICBM 2009c2 average template (at a resolution of 1 × 1 × 1 mm^3^) in addition to manual editing of cortical and subcortical labels.In the present dataset, we introduce an accurate non-linear registration of cortical and subcortical labelling from Mindboggle 101 to the symmetric MNI-ICBM2009c atlas followed by manual editing.

## Methods

### MNI-ICBM152 template

Within the ICBM project, MRI data from 152 young normal adults (18.5–43.5 years) were acquired on a Philips 1.5T Gyroscan (Best, Netherlands) scanner at the Montreal Neurological Institute (Mazziotta et al., 1995). The T1w data were acquired with a spoiled gradient echo sequence (sagittal acquisition, 140 contiguous 1 mm thick slices, TR=18 ms, TE=10 ms, flip angle 30°, rectangular FOV of 256 mm SI and 204 mm AP). The Ethics Committee of the Montreal Neurological Institute approved the study, and informed consent was obtained from all participants. ^2^

The following preprocessing steps were applied to all MRI scans prior to building the atlas: (1) N3 non-uniformity correction ^5^; (2) linear normalization of each scan’s intensity to the range [0-100] by a single linear histogram scaling ^6^; (3) automatic linear (nine parameters) registration to the ICBM 152 stereotaxic space^7^; and (4) brain mask creation^8^. Only the voxels within the brain volume after linear mapping into stereotaxic space were used for the nonlinear registration procedure described.

The template described is generated through a hierarchical nonlinear registration procedure, with diminishing step sizes in each iteration until convergence and relies on the nonlinear registration using Automatic Nonlinear Image Matching and Anatomical Labelling (ANIMAL)^9^. The nonlinear versions of MNI-ICBM2009 (http://nist.mni.mcgill.ca/?p=904) have many advantages over widely used previous versions (i.e. MNI-ICBM non-linear 6_th_ generation; http://nist.mni.mcgill.ca./?p=858 https://fsl.fmrib.ox.ac.uk/fsl/fslwiki/Atlases). Besides, anatomical variability still remains after linear transformation to stereotaxic space, therefore sulci and gyri remain blurred in previous versions^10^ (Figure 1)

Methodological details can be found in Fonov et al. 2011^2^.

**Figure 1.**
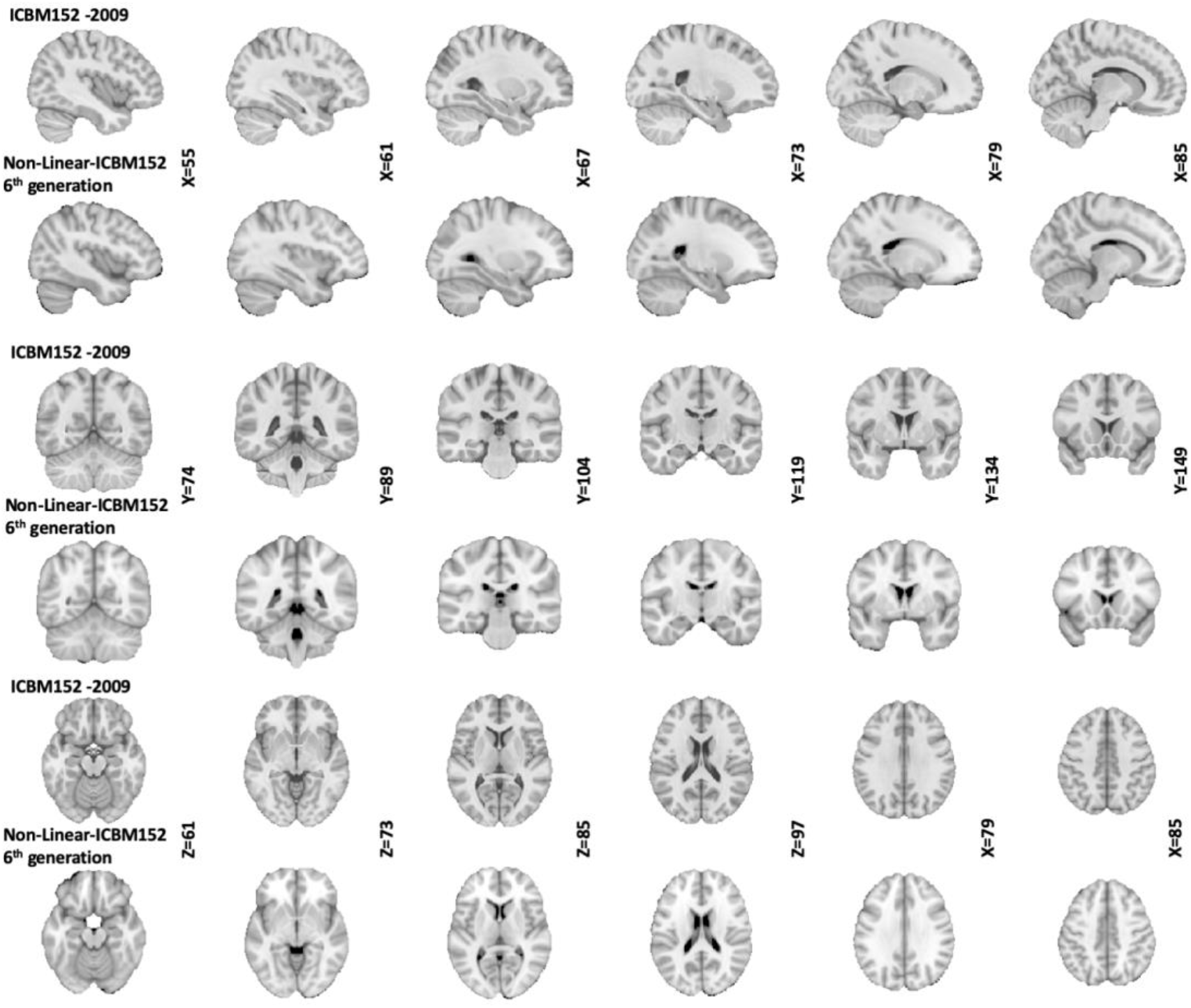
Comparison between non-linear 6^th^ generation MNI-ICBM and MNI-ICBM2009c

### Mindboggle-101

T1-w publicly accessible MRI scans were selected from 101 healthy participants. Scanner acquisition and demographic information can be found in Klein 2012^1^ and are also available on the http://mindboggle.info/data website. The data sets that comprise the Mindboggle-101 include the 20 test–retest subjects from the “Open Access Series of Imaging Studies” data^11^, the 21 test–retest subjects from the “Multi-Modal ReproducibilityResource”^12^, with two additional subjects run under the same protocol in 3 T and 7 T scanners, 20 subjects from the “Nathan Kline Institute Test–Retest” set, 22 subjects from the “Nathan Kline Institute/Rockland Sample”, the 12 “Human Language Network” subjects^13^, the Colin Holmes 27 templates, two identical twins, and one brain imaging colleague.

T1-w MRI volumes were preprocessed and segmented and then, cortical surfaces were generated using FreeSurfer’s standard recon-all image processing pipeline4^15,16^. FreeSurfer then automatically labelled the cortical surface using its DKT cortical parcellation atlas^3,17^. Vertices along the cortical surface are assigned a given label based on local surface curvature and average convexity, prior label probabilities, and neighbouring vertex labels. FreeSurfer automatically labelled the cortical surface using its DKT cortical parcellation atlas for 54 of the brains in the Mindboggle-101 data set. The region definitions of the labelling protocol represented by the DKT atlas are described by Desikan et al.^3^. These labels were then manually edited in agreement with the DKT protocol with 31 cortical regions per hemisphere as described by Klein and Tourville1. Then, the first 40 brains that labelled were selected to train a new FreeSurfer cortical parcellation atlas representing the DKT protocol (see http://surfer.nmr.mgh.harvard.edu/fswiki/FsTutorial/GcaFormat; S’egonne et al. 2004^17^; Desikan et al. 2006^3^ for details regarding the algorithm that generates the atlas and how it is implemented). The resulting atlas was named “DKT40 classifier atlas” which then automatically generated the initial set of cortical labels for the remaining 47 brains in the data.

Finally, Mindboggle data includes non-cortical labels that were converted from the Neuromorphometrics BrainCOLOR subcortex labels (i.e., http://Neuromorphometrics.com/^4^). Details on the original labels included in Mindboggle-101 can be found in https://mindboggle.readthedocs.io/en/latest/labels.html.

### Atlas registration and manual label editing

The Mindboggle-101 average template was non-linearly registered to the symmetric version of MNI–ICBM (MNI-ICBM2009c) template using the ANTs diffeomorphic registration pipelines^18^. Using the obtained nonlinear transformation, the Mindboggle-101 atlas labels were also resampled and registered to MNI-ICBM2009 template. The quality of the registration of the original labels was then visually assessed and inaccuracies were manually corrected on the right hemisphere using MNI Display (https://www.mcgill.ca/bic/software/visualization/display). Afterwards, labels were flipped onto the left hemisphere and then visual inspection on each structure was performed.

In detail, thickness and boundaries of all 51 cortical and subcortical labels from each hemisphere were improved using intensity thresholds with manual painting using MNI Display. Particular edits made for some structures are shown in Table 1.

**Table 1.** Original label numbers from Mindboggle with new label numbers. Table is showing specific corrections that were made to some structures for CerebrA and the agreement between the to labelling methods (Dice Kappa coefficient) Abbreviations: MB: Mindboggle-10; Vol: volume; MT: middle temporal; CMF: caudal middle frontal; IP: inferior parietal; CWM: cerebellar white matter; CSF: cerebrospinal fluid; OC: optic chiasm; CGM: cerebellar gray matter;

| MB ID | Label Name | Cerebra ID |  | Notes | Kappa |
| --- | --- | --- | --- | --- | --- |
|  |  | RH Labels | LH Labels |  |  |
| 2002 | Caudal Anterior Cingulate | 30 | 81 |  | 0.79 |
| 2003 | Caudal Middle Frontal | 42 | 93 | Improved distinction from Precentral | 0.73 |
| 2005 | Cuneus | 43 | 94 |  | 0.67 |
| 2006 | Entorhinal | 36 | 87 | Improved delimitation | 0.78 |
| 2007 | Fusiform | 24 | 75 |  | 0.77 |
| 2008 | Inferior Parietal | 10 | 61 |  | 0.75 |
| 2009 | Inferior temporal | 3 | 54 | Removed dorsal part MT | 0.72 |
| 2010 | Isthmus Cingulate | 33 | 84 |  | 0.79 |
| 2011 | Lateral Occipital | 34 | 85 |  | 0.76 |
| 2012 | Lateral Orbitofrontal | 7 | 58 |  | 0.8 |
| 2013 | Lingual | 12 | 63 |  | 0.75 |
| 2014 | Medial Orbitofrontal | 15 | 66 |  | 0.72 |
| 2015 | Middle Temporal | 28 | 79 | Added dorsal part | 0.72 |
| 2016 | Para hippocampal | 18 | 69 |  | 0.86 |
| 2017 | Paracentral | 16 | 67 |  | 0.77 |
| 2018 | Pars Opercularis | 32 | 83 |  | 0.77 |
| 2019 | Pars Orbitalis | 44 | 95 |  | 0.8 |
| 2020 | Pars Triangularis | 22 | 73 |  | 0.76 |
| 2021 | Pericalcarine | 6 | 57 |  | 0.6 |
| 2022 | Postcentral | 13 | 64 |  | 0.82 |
| 2023 | Posterior Cingulate | 47 | 98 |  | 0.8 |
| 2024 | Precentral | 35 | 86 |  | 0.84 |
| 2025 | Precuneus | 31 | 82 |  | 0.8 |
| 2026 | Rostral Anterior Cingulate | 8 | 59 |  | 0.72 |
| 2027 | Rostral Middle Frontal | 1 | 52 | Improved delimitation from CMF | 0.74 |
| 2028 | Superior Frontal | 38 | 89 |  | 0.82 |
| 2029 | Superior Parietal | 9 | 60 | Improved delimitation from Precuneus and IP | 0.72 |
| 2030 | Superior Temporal | 45 | 96 | Added dorsal part limiting with IP and Supramarginal | 0.87 |
| 2031 | Supramarginal | 51 | 102 |  | 0.81 |
| 2034 | Transverse Temporal | 14 | 65 |  | 0.85 |
| 2035 | Insula | 23 | 74 |  | 0.88 |
| 16 | Brainstem | 11 | 62 | Completed filling, removed labelled voxels out of actual brainstem and removed CWM labels in brainstem area. | 0.65 |
| 14 | Third Ventricle | 29 | 80 |  | 0.68 |
| 15 | Fourth Ventricle | 37 | 88 | Missing label. Manually delimited using CSF threshold. | 0.39 |
| 85 | Optic Chiasm | 17 | 68 | Almost inexistent label and out of place in original labelling Completed OC and tracts (originally labelled as Ventral Diencephalon) | 0 |
| 43 | Lateral Ventricle | 41 | 92 | Improved continuity of labelled voxels | 0.89 |
| 44 | Inferior Lateral Ventricle | 5 | 56 |  | 0.12 |
| 45 | Cerebellum Gray Matter | 46 | 97 | Completed filling using threshold for CGM, removed cerebellum labels out of area (within brainstem and vermis area) | 0.83 |
| 46 | Cerebellum White Matter | 39 | 90 | Improved according threshold for CWM, removed labels in brainstem and vermis. | 0.73 |
| 49 | Thalamus | 40 | 91 |  | 0.97 |
| 50 | Caudate | 49 | 100 | Completed filling using threshold | 0.84 |
| 51 | Putamen | 21 | 72 | Corrected uniformity using threshold | 0.87 |
| 52 | Pallidum | 27 | 78 | Improved delimitation between putamen and pallidum | 0.83 |
| 53 | Hippocampus | 48 | 99 |  | 0.69 |
| 54 | Amygdala | 19 | 70 |  | 0.64 |
| 58 | Accumbens Area | 4 | 55 |  | 0.76 |
| 60 | Ventral Diencephalon | 26 | 77 |  | 0.93 |
| 92 | Basal Forebrain | 25 | 76 |  | 0.82 |
| 630 | Vermal lobules I-V | 50 | 101 | Improved delimitation with other vermal lobules and cerebellar hemispheres | 0.66 |
| 631 | Vermal lobules VI-VII | 2 | 53 | Improved delimitation with other vermal lobules and cerebellar hemispheres | 0.38 |
| 632 | Vermal lobules VIII-X | 20 | 71 | Improved delimitation with other vermal lobules and cerebellar hemispheres | 0.44 |

## Data Records

CerebrA probabilistic atlas, including the corresponding T1w template, as well as segmentations of labels are available on http://nist.mni.mcgill.ca/?p=904. All imaging data are in compressed MINC format^19,20^. We invite contributions by other researchers, in terms of alternative opinions on labeling of included structures.

## Technical Validation

### Comparison between atlases

When comparing CerebrA to original labels from Mindboggle-101 (Figure 2) registered to ICBM152, the average Dice Kappa value was κ = 0.73 ± 0.18 (Table 1).

**Figure 2:**
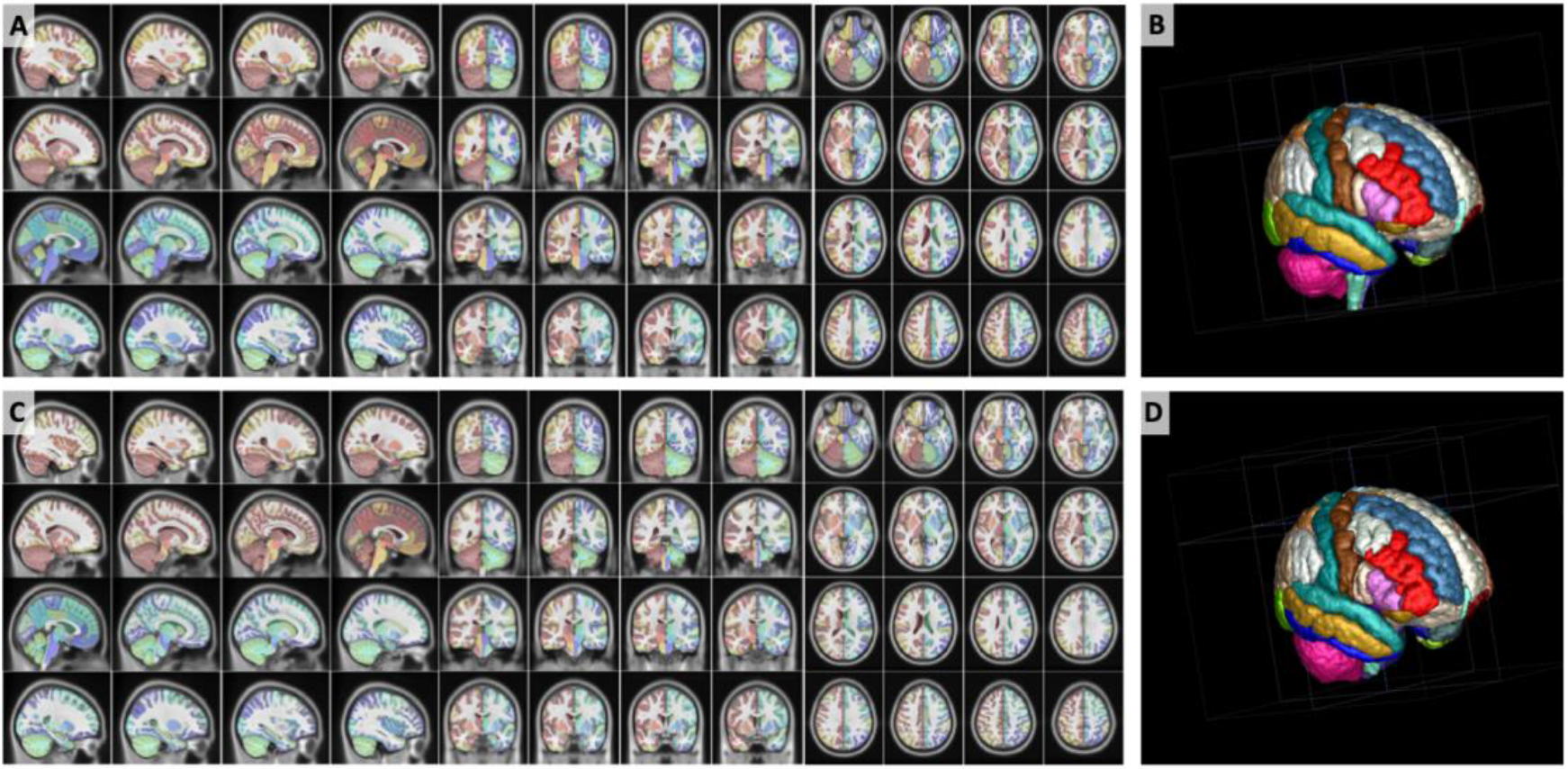
Warped CerebrA atlas a and b) and Mindboggle-101 atlas (c and d) overlaid on the ICBM152 non-linear 2009 symmetric average MRI template. Note the improved label alignment on the cortical structures.

The structures with relatively lower Dice Kappa (κ < 0.6) corresponded to the structures that needed the most correction such as the optic chiasm, inferior lateral ventricles, fourth ventricle and cerebellar vermis. The optic chiasm label was barely found in the original Mindboggle-101 registered to ICBM152 and most of it was misaligned with regards to the actual structure. To ensure that this inaccuracy was not caused by the nonlinear registration process, we further inspected the original Mindboggle-101 template and label atlas and found similar issues. For CerebrA, the optic chiasm label was redefined trying to achieve continuity amongst optic chiasma itself and optic tracts (Figure 3, panel a). Then, the inferior lateral ventricles and fourth ventricle boundaries were improved using a threshold to differentiate CSF from parenchyma (Figure 3, panels b and c). And finally, cerebellar vermis labels were redefined for right and left side (Figure 3, panels d-f).

**Figure 3.**
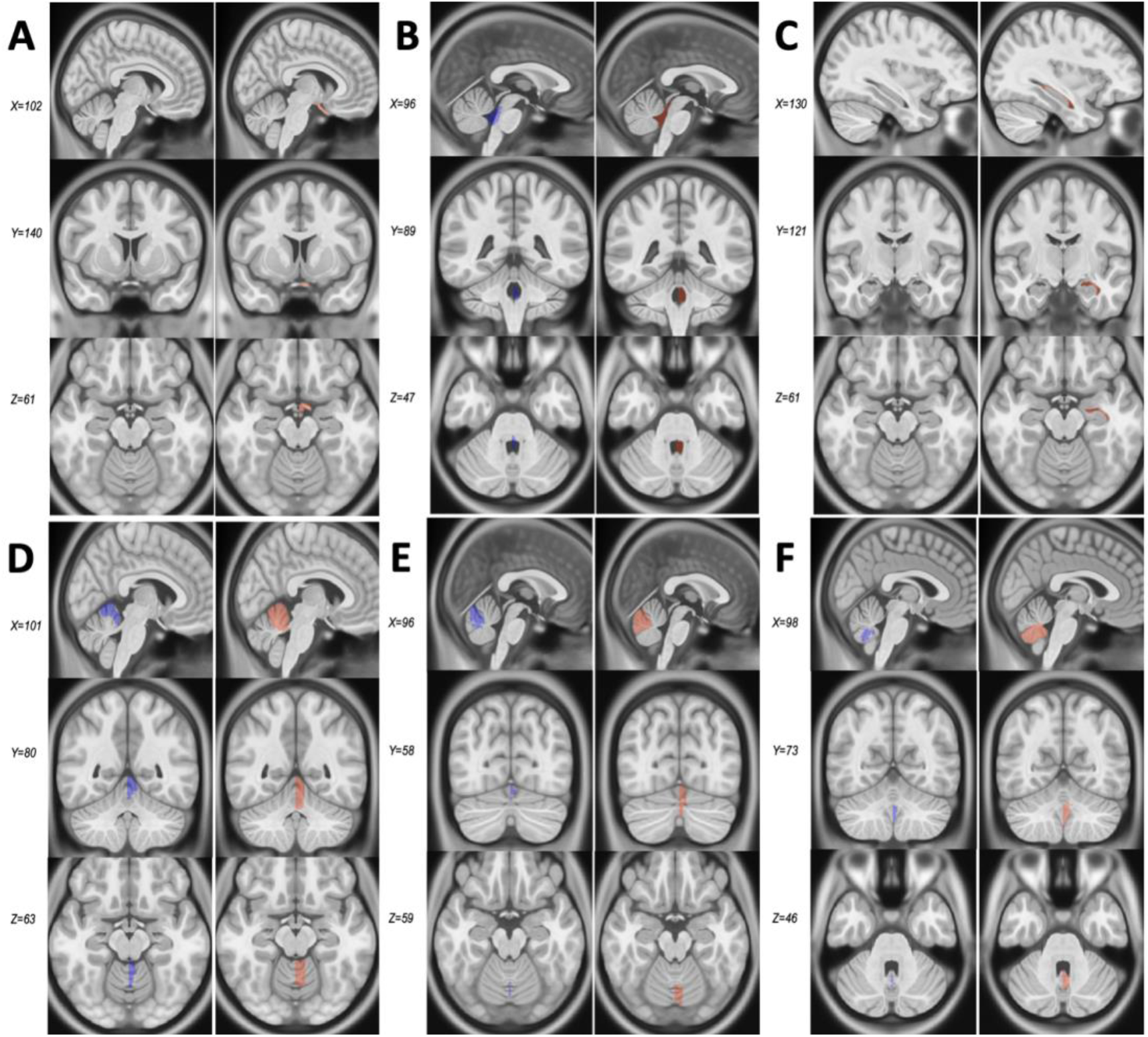
Comparison between Mindboggle-101 and CerebrA for structures with Dice Kappa < 0.6. **Panel a.** Optic chiasm. **Panel b.** Fourth ventricle. **Panel c.** Inferior lateral ventricle. **Panels d-f.** Cerebellar vermis lobules. For each structure the column on the left (purple) represents the original labels from Mindboggle-101, warped onto the ICBM152 symmetric template, and the right column (coral) represents CerebrA’s right sided corresponding labels, on the same template.

Another significant change in CerebrA from the original warped labels was the brainstem label definition. The brainstem area was manually redefined for the right side and then flipped in the same procedure as all the labels considering the symmetrical feature of the ICBM152 2009c^2^ template. In addition, boundaries between brainstem and fourth ventricle were carefully defined using the CSF intensity threshold, cerebellar white matter labels within the brainstem area were removed and rostral brainstem delimitation was improved (Figure 4).

**Figure 4.**
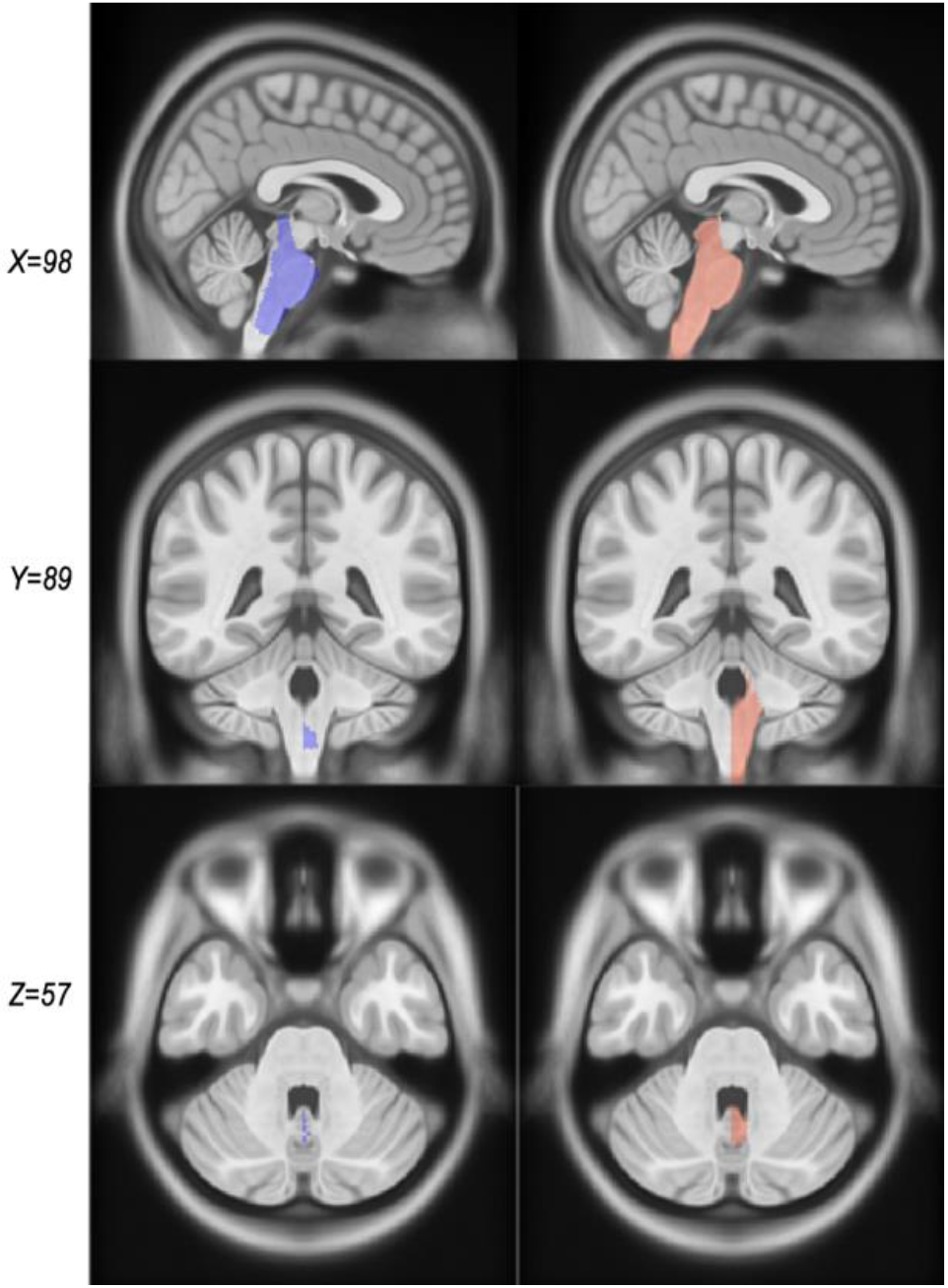
Comparison between warped Mindboggle-101 labels and CerebrA labels for the brainstem. The column on the left (purple) represents the original labels from Mindboggle-101 and the right column (coral) represents CerebrA’s right sided corresponding labels.

### Volumes of cortical and subcortical structures

In order to assess the size of the different structures, we calculated their volume by summing up the number of voxels with each label (in CCs). The volumes were then log-transformed to achieve normal distribution. Though significantly correlated (R= 0.9657, P value < 0.001), overall volumes estimated with CerebrA were larger than those estimated with Mindboggle-101 (Figure 5a). The volumes estimated per structure using Mindboggle and CerebrA segmentation are listed in Table 2.

**Figure 5.**
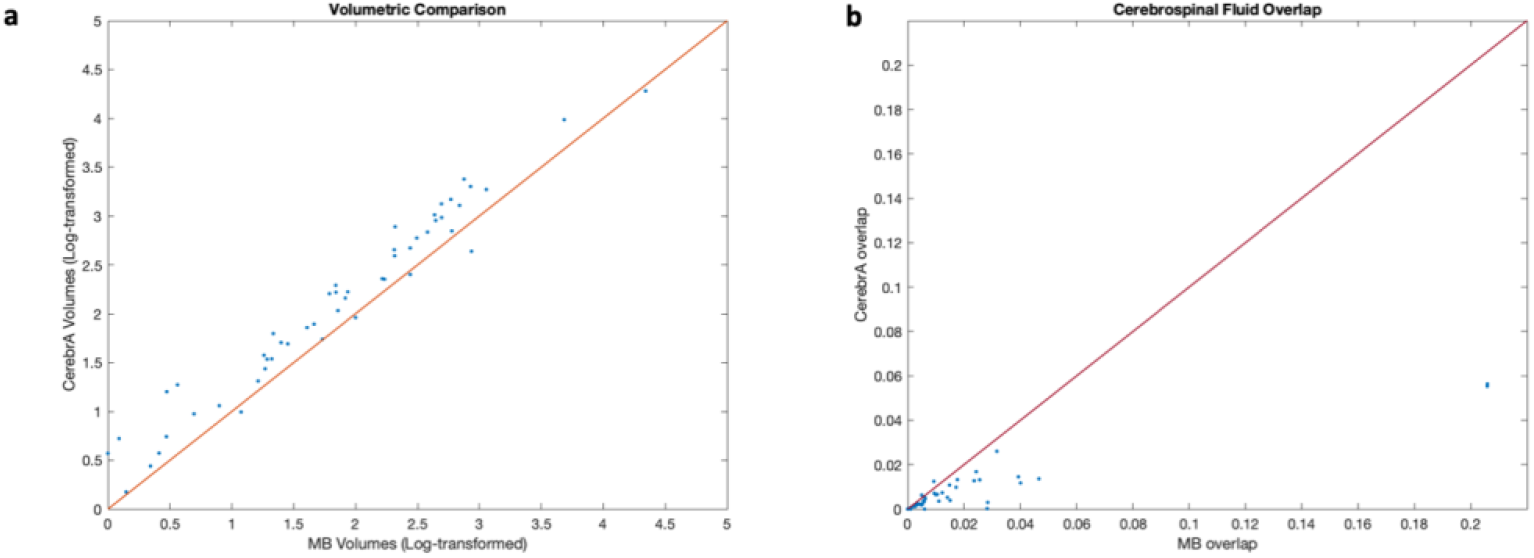
**a.** Correlation plot between volumes CerebrA and Minboggle-101. For better visualization and to achieve a normal distribution, the volumes have been log-transformed. Correlation coefficient R= 0.9657, P value < 0.001. **b.** Plot showing the proportion of overlap (the number of voxels in the specific mask overlapping with the CSF mask/the total number of voxels inside the specific mask) between gray matter labels and gray matter labels and cerebrospinal fluid for both atlases. Abbreviations: MB: Mindboggle-101

**Figure 6.**
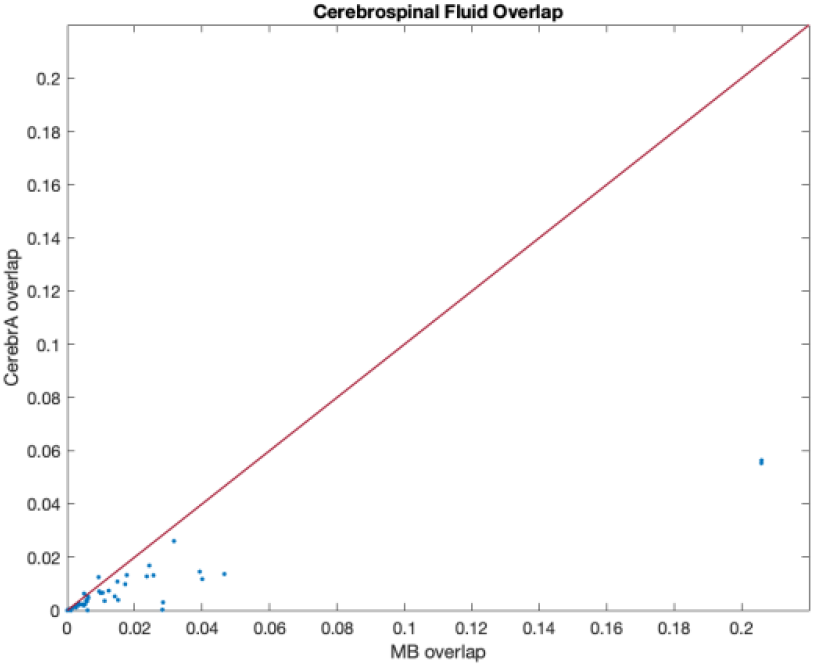
Plots showing the proportion of overlap (the number of voxels in the specific mask overlapping with the CSF mask/the total number of voxels inside the specific mask) between gray matter labels and gray matter labels and cerebrospinal fluid for both atlases. Abbreviations: MB: Mindboggle-101

**Table 2.** Volume per structure using Mindboggle-101 and CerebrA segmentations. Abbreviations: MB: Midboggle-101

| Label Name | Volume (cc) |  | Label Name | Volume (cc) |  |
| --- | --- | --- | --- | --- | --- |
|  | MB | CerebrA |  | MB | CerebrA |
| Caudal Anterior Cingulate | 2.75 | 3.65 | Superior Parietal | 12.97 | 19.33 |
| Caudal Middle Frontal | 9.1 | 13.22 | Superior Temporal | 20.25 | 25.35 |
| Cuneus | 4.97 | 8.07 | Supramarginal | 12.2 | 16.03 |
| Entorhinal | 2.56 | 3.21 | Transverse Temporal | 1.45 | 1.88 |
| Fusiform | 10.48 | 13.47 | Insula | 8.33 | 9.52 |
| Inferior Parietal | 17.73 | 26.19 | Brainstem | 9.17 | 17.01 |
| Inferior temporal | 15.1 | 16.22 | Third Ventricle | 0.6 | 1.1 |
| Isthmus Cingulate | 3.27 | 4.43 | Fourth Ventricle | 0.51 | 0.77 |
| Lateral Occipital | 14.97 | 22.81 | Optic Chiasm | 0 | 0.77 |
| Lateral Orbitofrontal | 11.1 | 15.04 | Lateral Ventricle | 8.16 | 9.57 |
| Lingual | 9.13 | 12.36 | Inferior Lateral Ventricle | 0.09 | 1.06 |
| Medial Orbitofrontal | 5.28 | 8.88 | Cerebellum Gray Matter | 75.85 | 71.23 |
| Middle Temporal | 16.77 | 28.29 | Cerebellum White Matter | 17.86 | 13 |
| Para hippocampal | 2.36 | 2.71 | Thalamus | 10.48 | 10.04 |
| Paracentral | 5.94 | 8.24 | Caudate | 4.28 | 5.64 |
| Pars Opercularis | 5.79 | 7.67 | Putamen | 5.4 | 6.63 |
| Pars Orbitalis | 2.61 | 3.64 | Pallidum | 1.93 | 1.7 |
| Pars Triangularis | 5.29 | 8.2 | Hippocampus | 4.64 | 4.69 |
| Pericalcarine | 2.8 | 5.03 | Amygdala | 1 | 1.65 |
| Postcentral | 13.1 | 18.19 | Accumbens Area | 0.41 | 0.55 |
| Posterior Cingulate | 3.99 | 5.41 | Ventral Diencephalon | 6.39 | 6.1 |
| Precentral | 16.13 | 21.39 | Basal Forebrain | 0.16 | 0.19 |
| Precuneus | 13.79 | 18.77 | Vermal lobules I-V | 3.05 | 4.5 |
| Rostral Anterior Cingulate | 2.52 | 3.83 | Vermal lobules VI-VII | 0.61 | 2.32 |
| Rostral Middle Frontal | 13.77 | 21.74 | Vermal lobules VIII-X | 0.75 | 2.57 |
| Superior Frontal | 38.87 | 52.82 |  |  |  |

### White matter and CSF overlap

Overall, Mindboggle-101 showed greater overlap of cortical and subcortical gray matter structures with cerebrospinal fluid (Figure 5b).

## Code Availability

The script used for generating the MNI-ICBM2009c average template is available at https://github.com/vfonov/build_average_model. Any other scripts used in the generation of the results of the current study are available from the corresponding author upon request.

## Authors contributions

**Ana L. Manera**: Study concept and design, manual correction of the labels, analysis and interpretation of the data, drafting and revision of the manuscript.

**Mahsa Dadar**: Study concept and design, analysis and interpretation of the data, revising the manuscript.

**Vladimir Fonov**: Generation of the MNI-ICBM2009c average template, revising the manuscript.

**D. Louis Collins**: Study concept and design, interpretation of the data, revising the manuscript.

## Acknowledgements

We would like to acknowledge funding from the Famille Louise & André Charron

## Abbreviations

MRI: magnetic resonance imaging
MNI: Montreal Neurological Institute
CerebrA: cerebrum atlas
DK: Desikan-Killiany cortical atlas
DKT: Desikan-Killiany-Tourville cortical atlas
T1w: T1-weighted

## References

1 Klein, A. & Tourville, J. 101 labeled brain images and a consistent human cortical labeling protocol. Front Neurosci 6, 171, doi:10.3389/fnins.2012.00171 (2012).

2 Fonov, V. et al. Unbiased average age-appropriate atlases for pediatric studies. Neuroimage 54, 313–327, doi:10.1016/j.neuroimage.2010.07.033 (2011).

3 Desikan, R. S. et al. An automated labeling system for subdividing the human cerebral cortex on MRI scans into gyral based regions of interest. Neuroimage 31, 968–980, doi:10.1016/j.neuroimage.2006.01.021 (2006).

4 Neuromorphometrics, I. <http://Neuromorphometrics.com/> (

5 Sled, J. G., Zijdenbos, A. P. & Evans, A. C. A nonparametric method for automatic correction of intensity nonuniformity in MRI data. IEEE Trans Med Imaging 17, 87–97, doi:10.1109/42.668698 (1998).

6 Nyul, L. G. & Udupa, J. K. On standardizing the MR image intensity scale. Magn Reson Med 42, 1072–1081, doi:10.1002/(sici)1522-2594(199912)42:6<1072::aid-mrm11>3.0.co;2-m (1999).

7 Collins, D. L., Neelin, P., Peters, T. M. & Evans, A. C. Automatic 3D intersubject registration of MR volumetric data in standardized Talairach space. J Comput Assist Tomogr 18, 192–205 (1994).

8 Smith, S. M. Fast robust automated brain extraction. Hum Brain Mapp 17, 143–155, doi:10.1002/hbm.10062 (2002).

9 A.C. Evans, D. L. C. a. Animal: validation and applications of nonlinear registration-based segmentation. International Journal ofPattern Recognition and Artificial Intelligence 11, 1271–1294, doi:10.1142/S0218001497000597 (1997).

10 Collins, M. P. D. L. in Atlas of the Morphology of the Human Cerebral Cortex on the Average MNI Brain (ed Academic Press) 17–22 (Natalie Farra, 2019).

11 Marcus, D. S. et al. Open Access Series of Imaging Studies (OASIS): cross-sectional MRI data in young, middle aged, nondemented, and demented older adults. J Cogn Neurosci 19, 1498–1507, doi:10.1162/jocn.2007.19.9.1498 (2007).

12 Landman, B. A. et al. Multi-parametric neuroimaging reproducibility: a 3-T resource study. Neuroimage 54, 2854–2866, doi:10.1016/j.neuroimage.2010.11.047 (2011).

13 Morgan, V. L., Mishra, A., Newton, A. T., Gore, J. C. & Ding, Z. Integrating functional and diffusion magnetic resonance imaging for analysis of structure-function relationship in the human language network. PLoS One 4, e6660, doi:10.1371/journal.pone.0006660 (2009).

14 Holmes, C. J. et al. Enhancement of MR images using registration for signal averaging. J Comput Assist Tomogr 22, 324–333, doi:10.1097/00004728-199803000-00032 (1998).

15 Dale, A. M., Fischl, B. & Sereno, M. I. Cortical surface-based analysis. I. Segmentation and surface reconstruction. Neuroimage 9, 179–194, doi:10.1006/nimg.1998.0395 (1999).

16 Fischl, B., Sereno, M. I. & Dale, A. M. Cortical surface-based analysis. II: Inflation, flattening, and a surface-based coordinate system. Neuroimage 9, 195–207, doi:10.1006/nimg.1998.0396 (1999).

17 Fischl, B. et al. Automatically parcellating the human cerebral cortex. Cereb Cortex 14, 11–22, doi:10.1093/cercor/bhg087 (2004).

18 Avants, B. B., Epstein, C. L., Grossman, M. & Gee, J. C. Symmetric diffeomorphic image registration with cross-correlation: evaluating automated labeling of elderly and neurodegenerative brain. Med Image Anal 12, 26–41, doi:10.1016/j.media.2007.06.004 (2008).

19 P. Neelin, J. M., D.L. Collins, and A. C. Evans. The minc file format: From bytes to brains (1998).

20 Vincent, R. D. et al. MINC 2.0: A Flexible Format for Multi-Modal Images. Front Neuroinform 10, 35, doi:10.3389/fninf.2016.00035 (2016).

